# Physical-biological interactions underlying the phylogeographic patterns of coral-dependent fishes around the Arabian Peninsula

**DOI:** 10.1101/2020.01.27.921411

**Authors:** Felipe Torquato, Peter R. Møller

## Abstract

**Aim:** To test the hypothesis that phylogeographic pattern of coral-dependent fish species inhabiting the Arabian Peninsula may be driven by a combination of ocean circulation, larval behavior and seascape features.

**Location:** The present study focuses on three such putative oceanographic barriers around the Arabian Peninsula: the Bab el-Mandeb Strait, the Strait of Hormuz and the upwelling off Oman.

**Tax:** Multitaxa.

**Methods:** A biophysical modeling system that relies on stochastic Lagrangian framework and Individual-Based Model was used to simulate larval dispersal through the three putative barriers, by tracking three-dimensional movements of virtual particles in ocean circulation scenarios. We explored the range of dispersal capabilities across reef fish species by creating 72 hypothetical strategies, each representing a unique combination of five biological traits: pelagic larval duration, spawning periodicity, mortality rate, reproductive output and vertical migration.

**Result:** Our results showed that the strength of the barriers was highly variable as a function of all biological traits (except reproductive output) and indicated high asymmetry of connectivity, and hence gene flow, between adjacent areas. In addition, direction and distance travelled by the virtual larvae varied according to both the geographic position of releasing site and biannual monsoonal winds. On average, larvae released during the summer exhibited a higher potential for dispersal than larvae released in wintertime.

**Main conclusions:** Our biophysical models complement the few existing empirical research on population genetics, and the predictions presented here serve as testable hypotheses for future phylogeographic studies around the Arabian Peninsula.

## INTRODUCTION

The Arabian Peninsula lies on a hyper-arid region in the Southwest Asia at the junction of this continent with Africa (Figure 1). The water-mass distribution and upper-ocean circulation surrounding the peninsula change in correspondence with biannual wind reversals, which creates seasonality in oceanographic conditions (Cutler & Swallow, 1984; Shetye et al., 1994). During the NE monsoon in the winter (November - March), the wind blows away from Asian continent, and the ocean surface circulation in the Arabian Sea is approximately counter-clockwise. On the other hand, during the summer SW monsoon (May-September) the wind reverses and blows strongly, so that the circulation in the Arabian Sea is clockwise. March-April and October are transition periods and winds are weak (see Cutler & Swallow, 1984).

**FIGURE 1.**
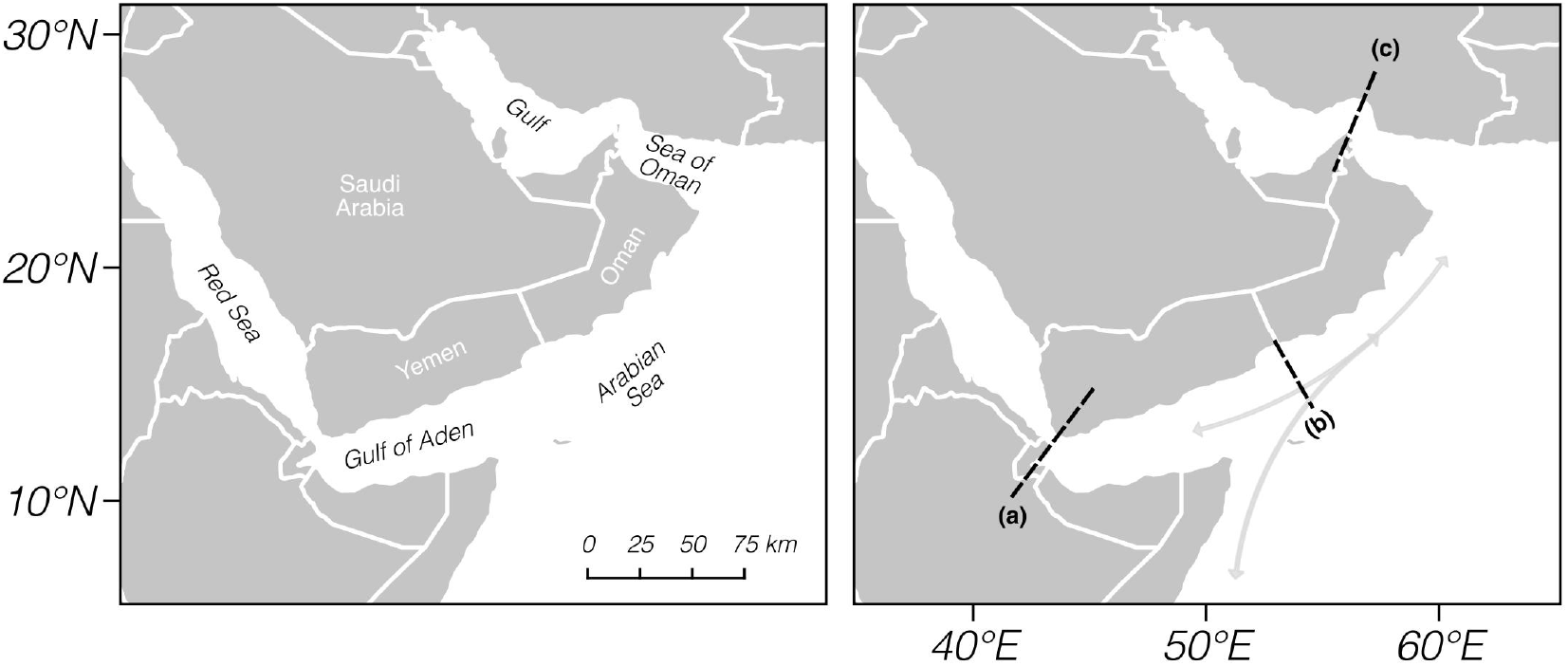
Study region. Northwestern Indian Ocean with previously described barriers depicted as shaded line in left panel. (a) Bab-el-Mandeb Strait, (b) Upwelling off Oman and (c) Strait of Hormuz. Arrows in the right panel represent prevailing currents in the northwestern Indian Ocean

Large seasonal variations of such magnitude, which is generally not found elsewhere (Shetye et al., 1994), plays a critical role for distribution of fish and other marine biodiversity by transporting or disrupting the movements of their larvae (Cowen, 2002). Thereby, the direction and magnitude of prevailing currents can spread out species distribution by aiding the dispersal of the propagules to future habitats. Nevertheless, these currents may also act as barriers by minimizing long-distance larval dispersal (Paris & Cowen, 2004), which ultimately may explain either the existence of endemic species (Cowen et al., 2000) or genetic discontinues across species ranges (Bowen et al., 2016). Although large expanses of ocean waters are the most notable permeable barriers for larval dispersal (Bowen et al., 2016), mesoscale oceanographic processes such as fronts (Galarza et al., 2009), river runoffs (Rocha, 2003), and upwellings (Lett et al., 2007) also acts as barriers to marine faunal connectivity. The permeability of these so-called semi-permeable barriers is taxon-specific and, therefore, affected by biological traits (Ayre et al., 2009).

At the Arabian Peninsula, zoogeographic and population genetic studies on coral-dependent fishes have shown discontinuities in both species and genetic distributions along the seas surrounding the peninsula (Burt et al., 2011; DiBattista, Roberts, et al., 2016; Berumen et al., 2017). The most remarkable of these discontinuities, distinguishes the fish fauna from each side of the peninsula (DiBattista, Choat, et al., 2016). Two other discontinuities in species compositions are observed, one between the Red Sea and the adjacent Gulf of Aden through the Bab el-Mandeb Strait, and another between the Arabian/Persian Gulf (henceforth referred as the Gulf) and the adjacent Sea of Oman through the Strait of Hormuz (DiBattista, Roberts, et al., 2016). These boundaries described by multispecies distribution records, are also logical places in relation to present-day barriers for gene flow between populations (Baums et al., 2006). Indeed, population genetic studies carried out in the Northwestern Indian Ocean (NIO) have revealed similarities in the geographical position of barriers previously proposed (DiBattista et al., 2013, 2015, 2017; Nanninga et al., 2014; Priest et al., 2016; Saenz-Agudelo et al., 2015; Torquato et al., 2019).

Hypotheses to explain the distribution of genetic diversity in the Arabian Peninsula usually rely on parapatric speciation pattern, which resulted from either repeated vicariance events caused by lowering sea level during the last glaciation, or ecological speciation due to the large spatial gradients and temporal fluctuations in physical conditions across the peninsula (DiBattista, Roberts, et al., 2016; DiBattista, Choat, et al., 2016; Nanninga et al., 2014). An alternative hypothesis has been attributed to the seasonal upwelling in the Arabian Sea off Oman, which creates unsuitable condition for discrete coral-habitat growth along southern Omani coast (Sheppard & Salm, 1988) and hence potentially restrict stepping-stone connectivity between both sides of the Arabian Peninsula. In turn, little attention has been given to test hypotheses where the combination of seascape features, ocean circulation and larval traits underling the genetic patterns observed among coral-dependent fishes inhabiting the Arabian Peninsula.

In numerous cases, hypotheses in marine ecology are prohibitively time consuming and expensive to be empirically tested, largely owing to the impossibility of capturing the full range of temporal and spatial fine-scale resolution required to make inferences (Cowen & Sponaugle, 2009). Although a more comprehensive picture is emerging in the Arabian Peninsula with respect to marine phylogeographic and population genetic patterns (Berumen et al., 2017; DiBattista, Roberts, et al., 2016; DiBattista, Choat, et al., 2016), the processes affecting larval dispersal across the putative barriers are not yet fully understood due to the paucity of empirical studies. In addition, political realities of some countries bordering the Western Indian Ocean (WIO) have limited access to scientists, hindering a more complete perspective of general phylogeography in the region (Berumen et al., 2017). In such cases, if limited empirical data are available, reliable computational models can be used to make field predictions and advance our knowledge to designing future experiments for hypothesis testing (Cowen & Sponaugle, 2009).

Advances on physical circulation models enabled the investigation of population connectivity by running semi-realistic simulations of virtual particles. Here we used high-resolution ocean circulation model (Hybrid Coordinate Ocean Model - HYCOM) to design a biophysical model in a Lagrangian stochastic scheme (Paris et al., 2013). The main goal of this study is to simulate multitaxon larval dispersal through the marine barriers, and thus provide insights on processes and patterns of connectivity leading to the distribution of genetic diversity of coral-dependent fishes around the Arabian Peninsula. Specifically, we shed light on three questions: (1) what biological attributes affect the larval dispersal through the Bab el-Mandeb Strait and Strait of Hormuz? (2) What is impact of the upwelling off Oman on the larval connectivity between both sides of the peninsula? (3) How does oceanographic variability, due to the seasonal monsoon, affect larval dispersal pattern (i.e. direction and magnitude travelled by the particle)? Our results provide detailed predictions that can be compared to previous and future empirical studies on the distribution of biodiversity in the Arabian Peninsula.

## MATERIAL AND METHODS

### Biophysical Model and Larval Dispersal Simulation

Idealized dispersal of fish larvae is modeled using an open-source program, Connectivity Modelling System (CMS v. 2.0; Paris et al., 2013), which is a biophysical modeling system based on stochastic Lagrangian framework and Individual-Based Model (IBM) that couple ocean current, GIS-based habitat, and biological traits. In brief, CMS uses information on currents and environmental conditions to simulate both deterministic fourth-order Runge-Kutta and/or stochastic displacements of a large number of virtual particles (hereafter called larvae), through space and time. In order to explore the range of dispersal capabilities across reef fishes, we created 72 hypothetical strategies each of those representing a unique combination of five biological traits that may influence the connectivity: pelagic larval duration (PLD), spawning periodicity, mortality rate, reproductive output and vertical migration (Table 1).

**TABLE 1.** Range in biological parameter values to characterize the 72 hypothetical model taxa.

| PLD (days) | Spawning periodicity | Larval mortality | Vertical migration | Reproductive output |
| --- | --- | --- | --- | --- |
| 20 | Annual | High | Yes | High |
| 30 | Summer) | Low | No | Low |
| 40 | Winter) |  |  |  |

### Hydrodynamic model

We use the CMS package *getdata* to download daily ocean current velocities from the three-dimensional and eddy-resolving Hybrid Coordinate Ocean Model (HYCOM, GLBa0.08) from 2014 to 2016. The model has a horizontal resolution of ca. 9 km grid (1/12°), and is set up in a nested domain (0° - 30° N, 31° - 70° E), comprising all layers from the surface up to 100 meters depth.

### GIS-based Seascape module

The GIS serves to delineate the source and recruitment habitats. Suitable releasing (source habitat) and settlement (sink habitat) locations are delineated in QGIS v.2.18 by creating a vector grid that overlaid the distribution of coral reefs data from the UNEP-WCMC (2010). A total of 181 polygons (∼ 18km x 18km) representing coral reef habitats are placed along coastal areas within our model domain.

### Biological module

We assume spatial homogenous reproductive output across the habitats and, therefore, all the 181 polygons are set to release from their center the same number of larvae per ‘spawning’ event. However, we contrast scenarios where reproductive output varies among species. In scenarios of high reproductive output, a hundred larvae are released from each polygon, whereas in low reproductive output conditions only ten are released. Here, three seasonal spawning preferences are accounted by simulating the larval dispersal annually according to the two dominant seasonal oceanographic conditions around the Arabian Peninsula. After being released, the larvae are let to drift over a period of ^20, 30^ or 40 days, corresponding to the typical pelagic larvae duration (PLD) of most coral reef fish species/families (Lindeman et al., 2005; Thresher and Brothers 1985).

In this study the PLD is equally divided into the three ontogenetic larval stages, namely: preflexion, flexion and postflexion. Due to the paucity of basic data regarding fish larvae distribution within our study area, the model incorporates an idealized pattern of ontogenetic vertical migration for the three larval stages (see Appendix S1 in Supporting Information). This idealization assumes the existence of a global ontogenetic trend of the fish larvae to display downward ontogenetic shift in vertical distribution (Irisson et al., 2010). During the postflexion stage the larvae are considered competent to settle if they are inside one of the 181 reef sites. In order to assess the importance of vertical migration, which potentially allow the larvae to avoid passive advection in vertically stratified scenarios and increase the chance of retention (Paris & Cowen, 2004), we also contrasted both epipelagic ichthyoplankton moving only horizontally along the sub-surface layer and larvae that, besides the horizontal displacement, also move vertically according to its ontogeny (Figure S1).

Little is known regarding larval mortality in the ocean. To accommodate this uncertainty, our study includes two levels of mortality based on the half-life, such that approximately 50% of unsettled larvae would be surviving after half the maximum PLD (Holstein et al., 2014; Paris et al., 2013). Thus, we determines that in high mortality scenario about half of the larvae die by the end of the preflexion stage, while in low mortality condition half of the larvae dies by the end of the postflexion stage (Table 1).

## Particle-tracking module

Stochastic IBM Lagrangian model tracks offline over 157 millions larvae around the Arabian Peninsula for the 72 hypothetical strategy. A total of 86,011,200 larvae are released to mimic taxa spawning throughout year, whereas 35,838,000 larvae are tracked to simulate taxa spawning on either winter monsoon (November 2014 – March 2015) or summer monsoon (May 2015 – September 2015).

Preliminary sensitive analysis showed no significant difference in the settlement proportion when seeding 100 larvae from the 181 polygons at every either 3 or 24 hours (see Appendix S1). Therefore, the larvae are released from each reef at every 24 hours, which represents a spawning event. This uniform temporal distribution of larvae allows us to assess the effects of the hydrographic variability conditions on larval dispersal (e.g., extreme events, perturbation, and instability). Throughout the PLD, the position of each larva is updated every 6 h time-step, and the trajectory information (i.e. longitude, latitude, depth) is saved to output in intervals of 24 hours. We account for diffusive turbulent motion by adding a horizontal diffusion coefficient. The value of 50 m^2^.s^-1^ was chosen from a sensitivity test where 100 larvae were released from the 181 polygons at every 24 hours (see Appendix S1).

### Analyses

The proportion of survivor larvae that were released from each region *i* and successfully settled in a downstream habitat patch (i.e., sink habitat) at region *j*, is plotted as a connectivity matrix. Self-recruitment, i.e. larvae settling within its release location, is represented by the diagonal of the matrix. In order to evaluate the strength and direction of the potential connections on regional scale (e.g., between Red Sea and Gulf of Aden), all cells from each region are merged.

### Biological traits vs. biogeographic barriers

To evaluate the effect of the biological traits highlighted in Table 1 on the putative barriers, we rely on a multiple regression approach. Specifically, the effects of both Bab-el-Mandeb and Hormuz strait are measured in terms of permeability, which considers the proportion of surviving larvae that are released from a source habitat, passed through the strait, and successfully settled on the other side. In turn, the effect of the upwelling off Oman is measured in terms of self-recruitment proportion along the Arabian Sea. At the region off Oman, we hypothesized that higher self-recruitment represent higher retention of larvae on continental shelf, and thus greater chance of connectivity. By contrast, we assume that lower self-recruitment is due to larval movement toward offshore as a consequence of Ekman transport, hence decreasing the change of the larvae to find suitable habitat for settlement (but see Morgan et al., 2012).

Provided that the proportion is a continuous variable that can take on values restricted to the interval between 0 and 1, a beta regression model as proposed by Ferrari & Cribari-Neto (2004) is used through the *betareg* R-package (Cribari-Neto & Zeilis, 2010). The beta regression is essentially similar to a Generalized Linear Model (GLM), where it describes the relationship between the response variable Y _i_(hereby, proportions) and the predictors X_i_(Table 1) through a linear predictor η_i_. This linear predictor is then linked to the mean of the response E(Y_i_)=µ_i_ by means of a link function g, such that g(µ_i_) = η_i_. In this way, the applied model can be summarized as:

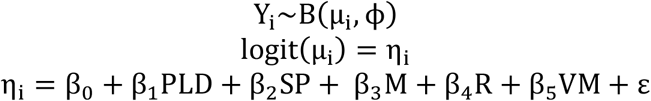

Where Φis the precision parameter; η_i_the linear predictor expressed on a logit scale; β0 the intercept; β1-5 the repressors related to each explanatory variable (PLD=Pelagic larval duration, SP = Spawning periodicity, M = Mortality, R = Reproductive output, VM = Vertical migration); and εrepresents the error term. Model assumptions like residual homocedasticity, independence and normality are accessed through conventional graphical checks as mentioned in Zuur et al. (2010).

### Larval dispersal pattern and oceanographic conditions

The maximum distances travelled by the larvae around the Arabian Peninsula are calculated by setting up a model with the lowest mortality rate and 40-days PLD. We use the program QGIS v.2.18 to measure this distance and compare the results in both summer and winter, as well as in each of the regions within the model domain.

## RESULTS

### Connectivity matrix

All the matrices representing the 72 hypothetical strategies revealed that connectivity occurred exclusively between neighbor regions (Figure S2 – S4). The hardest putative barrier was observed off Oman, which separates both sides of the Arabian Peninsula. Thereby, larvae that were released from the Arabian Sea, the Sea of Oman or the Gulf, rarely reached the reefs in the Gulf of Aden or the Red Sea, and vice-versa.

In turn, the connectivity through both the Bab el-Mandeb Strait and the Strait of Hommuz were not symmetric. Instead, the strength and symmetry of the connections varied either seasonally or due to larval traits. For example, for 7 strategies in which the larvae were released from the Red Sea, the larvae did not reach the reefs located in the adjacent Gulf of Aden. Likewise, for 5 strategies where larvae where released from the Sea of Oman, the larvae were not able to cross the Strait of Hommuz and settle onto reefs in the Gulf. In the former example, the thorough larval retention in the Red Sea occurred only in the winter and irrespective of the PLD. The second example, on the other hand, was only observed for the shortest PLD but irrespective of seasonality.

### Effects of the biological traits on the permeability of biogeographic barriers

Graphical evaluation of the model assumptions revealed that all tested models fitted well to the data, with residuals following both homoscedasticity and normality assumptions. The numerical outputs of the models are summarized in table S1 and discussed in more detail below.

### Bab el-Mandeb Strait

For the larvae that were released from the Red Sea and successfully settled in the Gulf of Aden, only the larval reproductive output was not significant (Table S1, Figure 2). The beta regression showed that the connectivity in this direction was significantly higher during the summer than winter, such that larvae experiencing high mortality in winter and that did not performed vertical migration, could not settle on the Gulf of Aden (Figure S2-S4). The permeability of the barrier in this direction was higher for larvae that performed vertical migration and were left to drift for longer periods, though no significant difference was observed between larvae that drift for 30 and 40 days.

**FIGURE 2.**
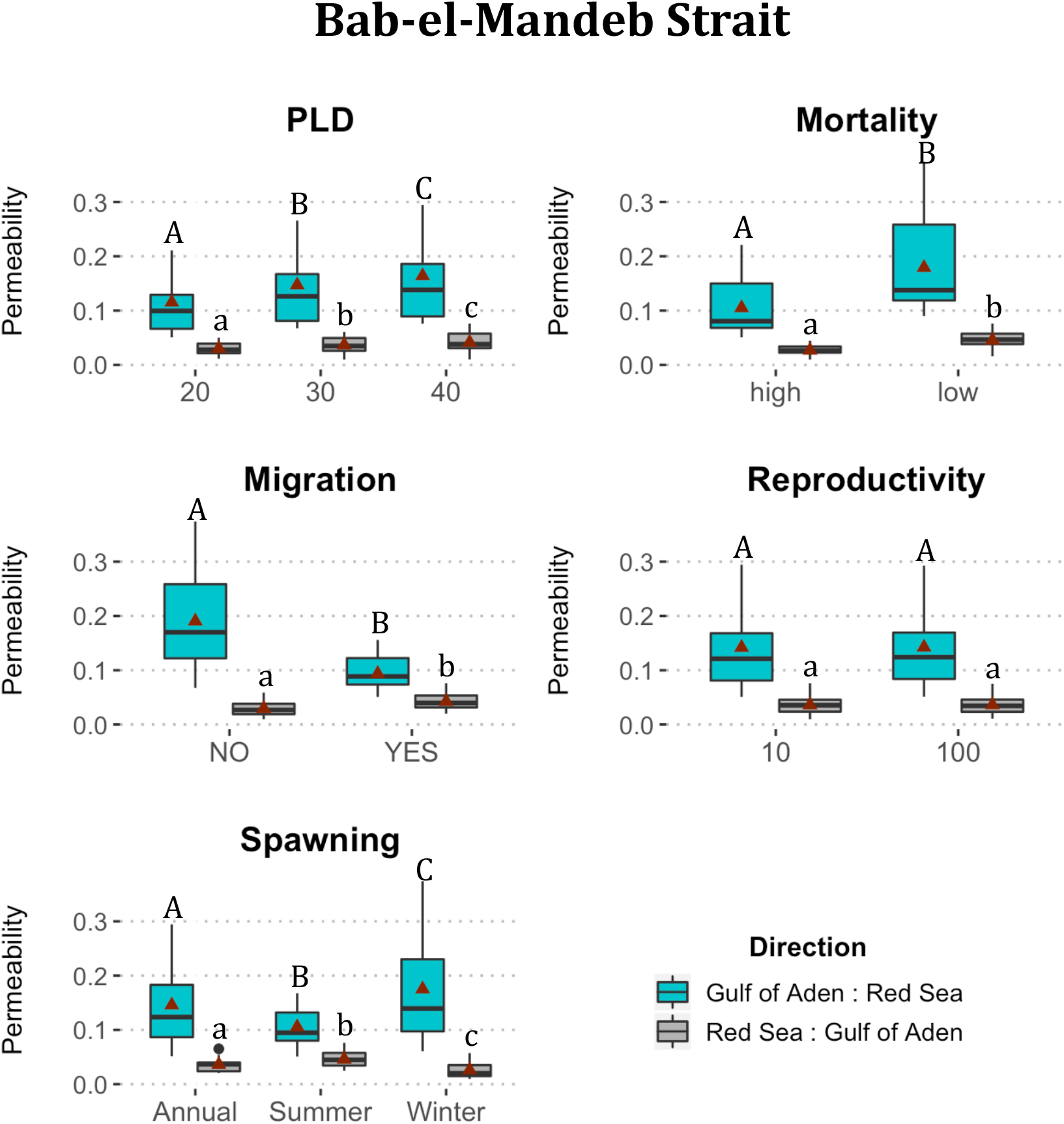
Results on the beta regression models showing the effects of the biological traits on the permeability of the Bab-el-Mandeb Strait. The red triangles represent the mean value. Capital and lowercase letters distinguish the direction showed in the legend, while differences between letters (a, b and c) indicate significant differences (p < 0·05).

On the other hand, larvae moving in the opposite direction, from the Gulf of Aden to the Red Sea, always crossed the Bab el-Mandeb Strait and settled on the other side. The higher success in the connection occurred during the wintertime, and although the larval reproductive output did not change the barrier’s permeability (Table S1, Figure 2), a positive relationship was observed between the permeability and higher PLD and no vertical migration. Mortality rate was also significant and displayed the same tendency of the larvae moving from the Red Sea and to the Gulf of Aden.

### Strait of Hormuz

The larvae release from the Gulf always reached the Sea of Oman. The proportion of larvae settling released in the second increased when they were released throughout the year or during the wintertime, performed vertical migration, had low mortality rate, and travelled for longer period (Table S1, Figure 3). Nevertheless, the chance of the larvae to cross the strait in this direction did not depend on the reproductive output.

**FIGURE 3.**
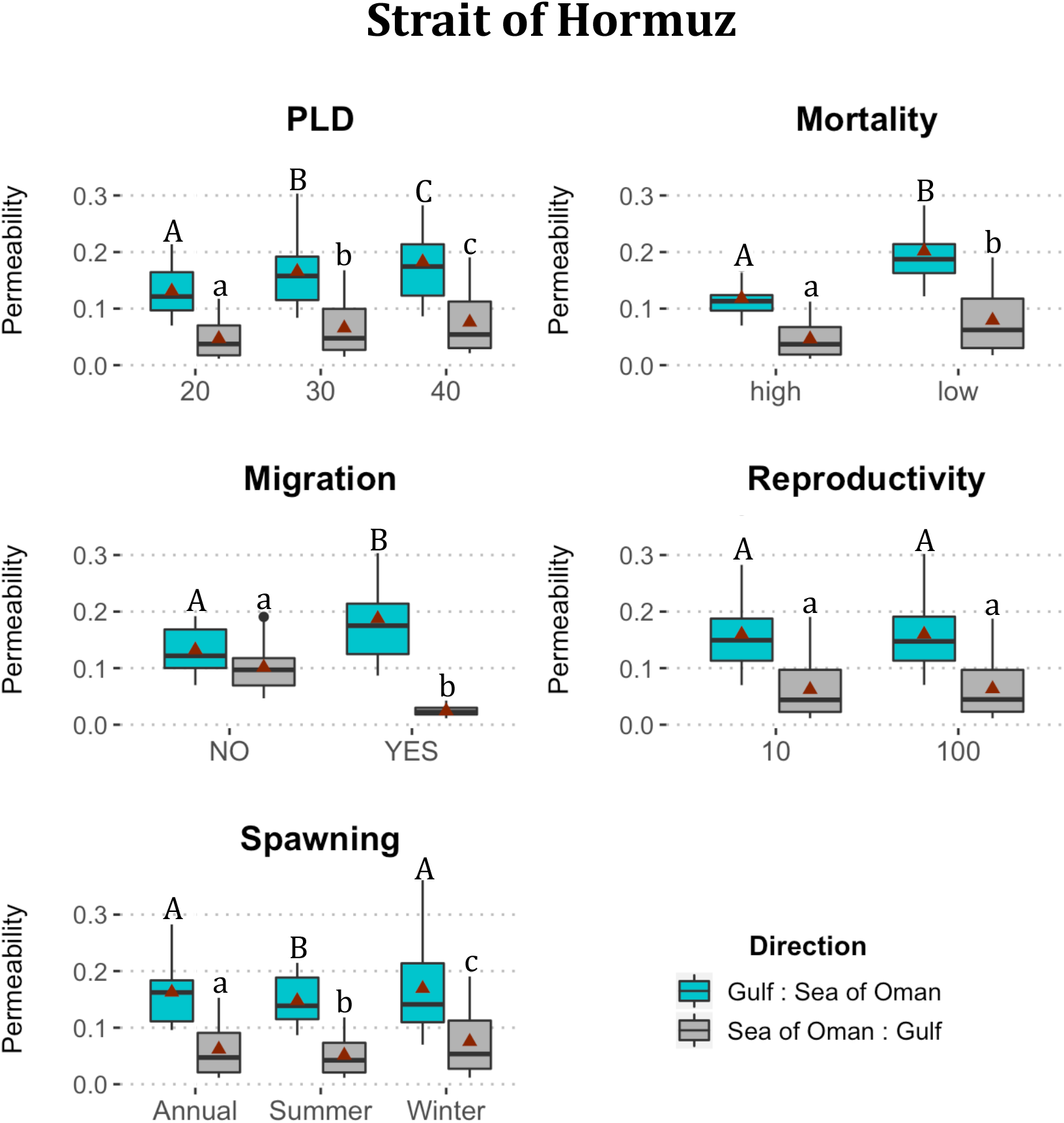
Results on the beta regression models showing the effects of the biological traits on the permeability of the Strait of Hormuz. The red triangles represent the mean value. Capital and lowercase letters distinguish the direction showed in the legend, while differences between letters (a, b and c) indicate significant differences (p < 0·05).

Likewise, among all biological attributes, only reproductive output was not significant for larvae that were released from the Sea of Oman, crossed the Strait of Hormuz and successfully settled in the Gulf. The probability of this connectivity was higher in the scenarios where the larvae were release in winter, had low mortality rate, did not perform vertical migration and remained in the planktonic environment for longer period. (Table S1, Figure 3). Importantly, 20-days PLD larvae, presenting high mortality and that performed vertical migration, did not cross the Strait of Hormuz (Figure S2).

### Upwelling off Oman

All biological traits, except larval reproductive output, significantly affected the proportion of self-recruitment in the Arabian Sea (Table S1, Figure 4). The scenario where the strategy had higher success in self-recruitment, and hence higher potential for connectivity, occurred during the wintertime or throughout year, and if the 20-days PLD larvae performed ontogenetic migration under low mortality scenario. In turn, larvae produced during the summer, when the Ekman transport takes place, had a slightly higher chance to be displaced to areas of unsuitable habitats.

**FIGURE 4.**
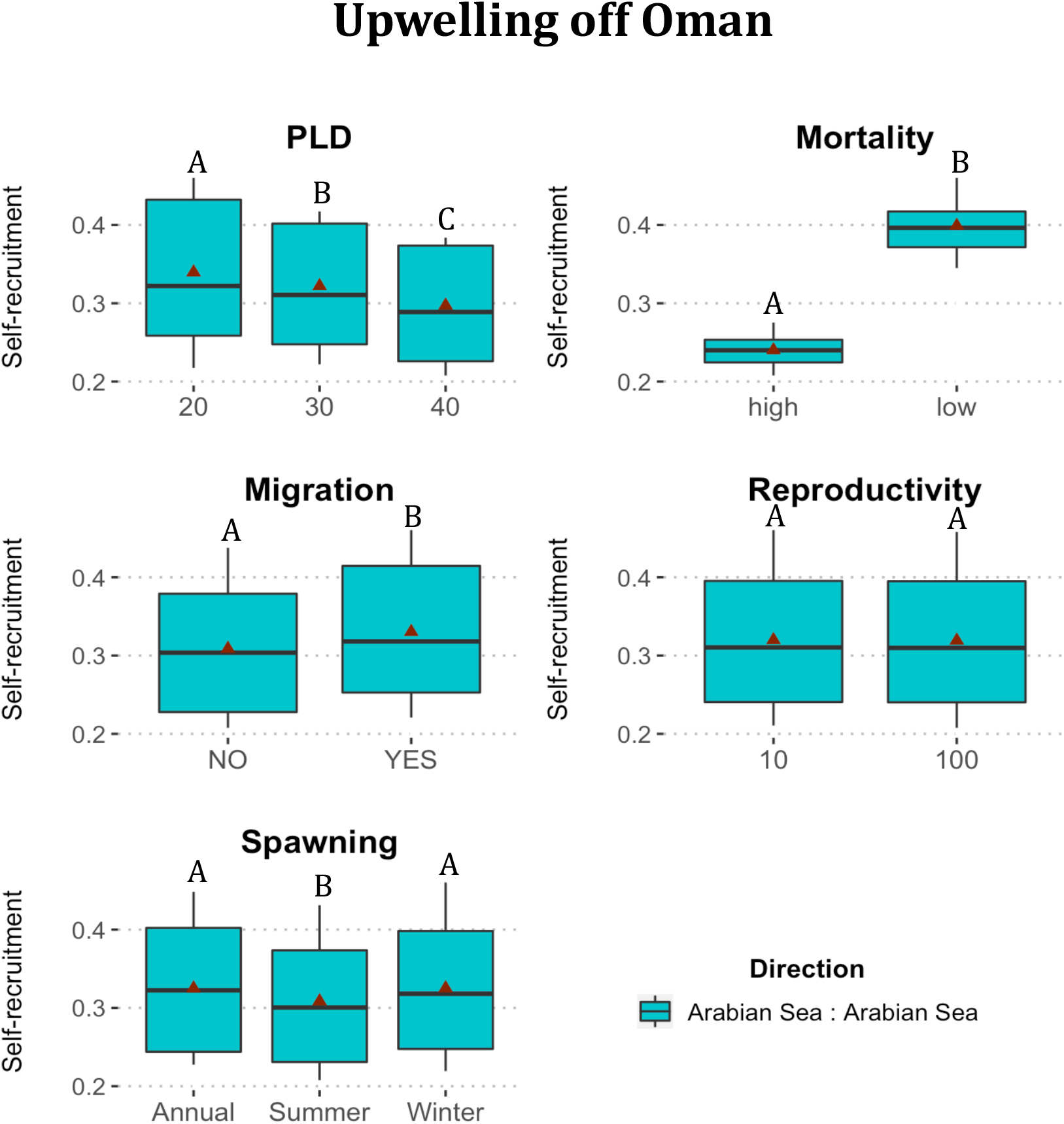
Results on the beta regression models showing the effects of the biological traits on the self-recruitment along Omani coast. The red triangles represent the mean value. Differences between letters (a, b and c) indicate significant differences (p < 0·05).

### Particles trajectories: seasonal variability of spatial scale and direction

The direction and distance travelled by the larvae were highly variable as a function of both the geographic position of releasing site and the biannual monsoonal winds (Figure 5 and 6). On average, larvae released during the summer exhibited a higher potential for dispersal, especially in areas that are not enclosed (Figure 5f, 6f). It was in summer that the movement of larvae southward from the Red Sea was more pronounced, and it was also in this season that many larvae released in the Gulf of Aden reached Omani waters in the Arabian Sea. In turn, during the winter the vast majority of larvae originating in the Red Sea were retained within this area, while larvae released within the Gulf of Aden either moved eastward, though rarely reaching the Omani coast, or moved southward along the Somali coast.

**FIGURE 5.**
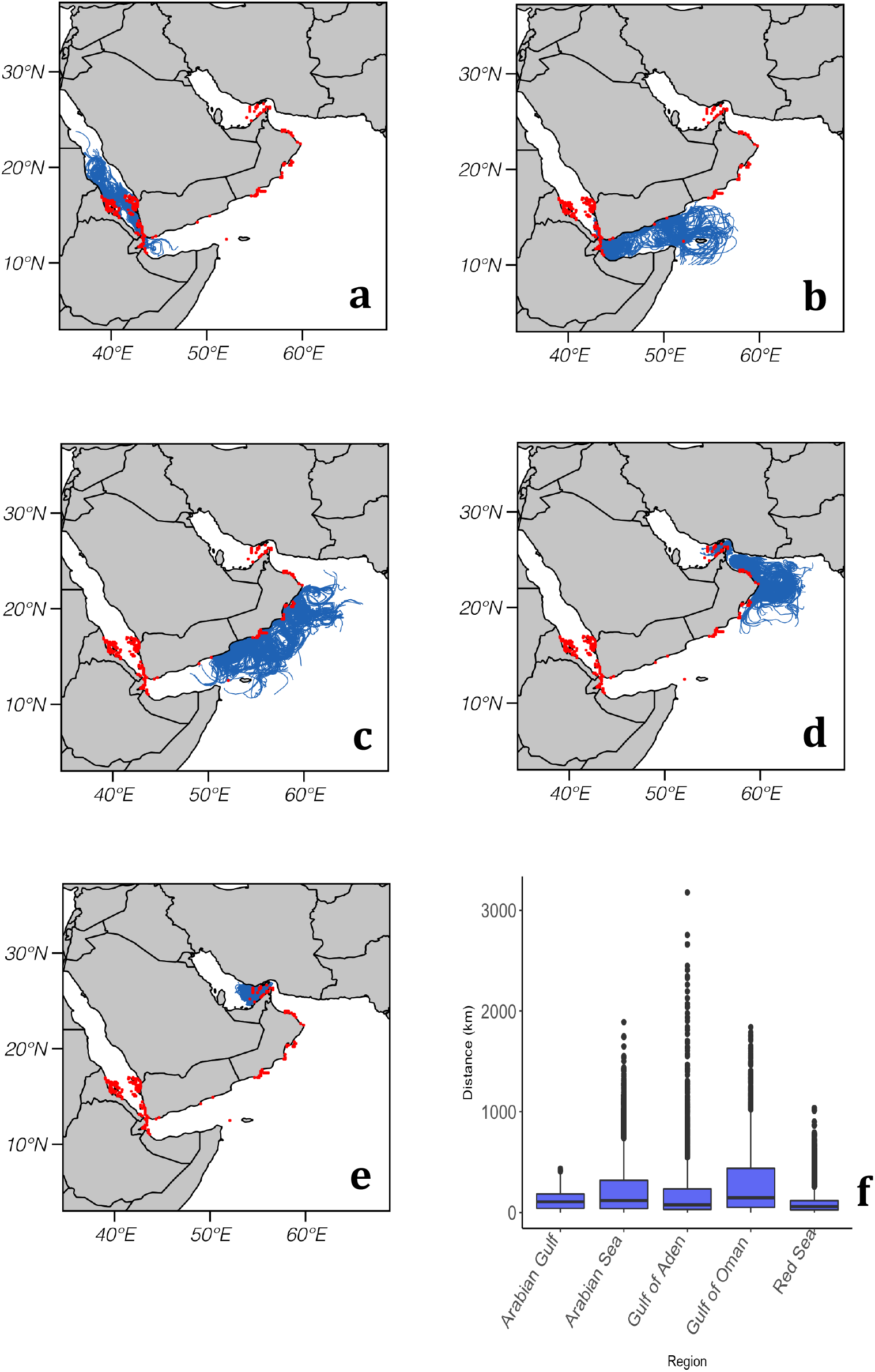
Modeled dispersal paths of epipelagic virtual larvae with larval duration of 40 days released in the wintertime from (a) the Red Sea, (b) Gulf of Aden, (c) Arabian Sea, (d) Sea of Oman and (e) Arabian Gulf. For sake of visualization one larva was released from each habitat cell (total of 181) per day in 2015. The distances travelled by the larvae are represented in (f).

**FIGURE 6.**
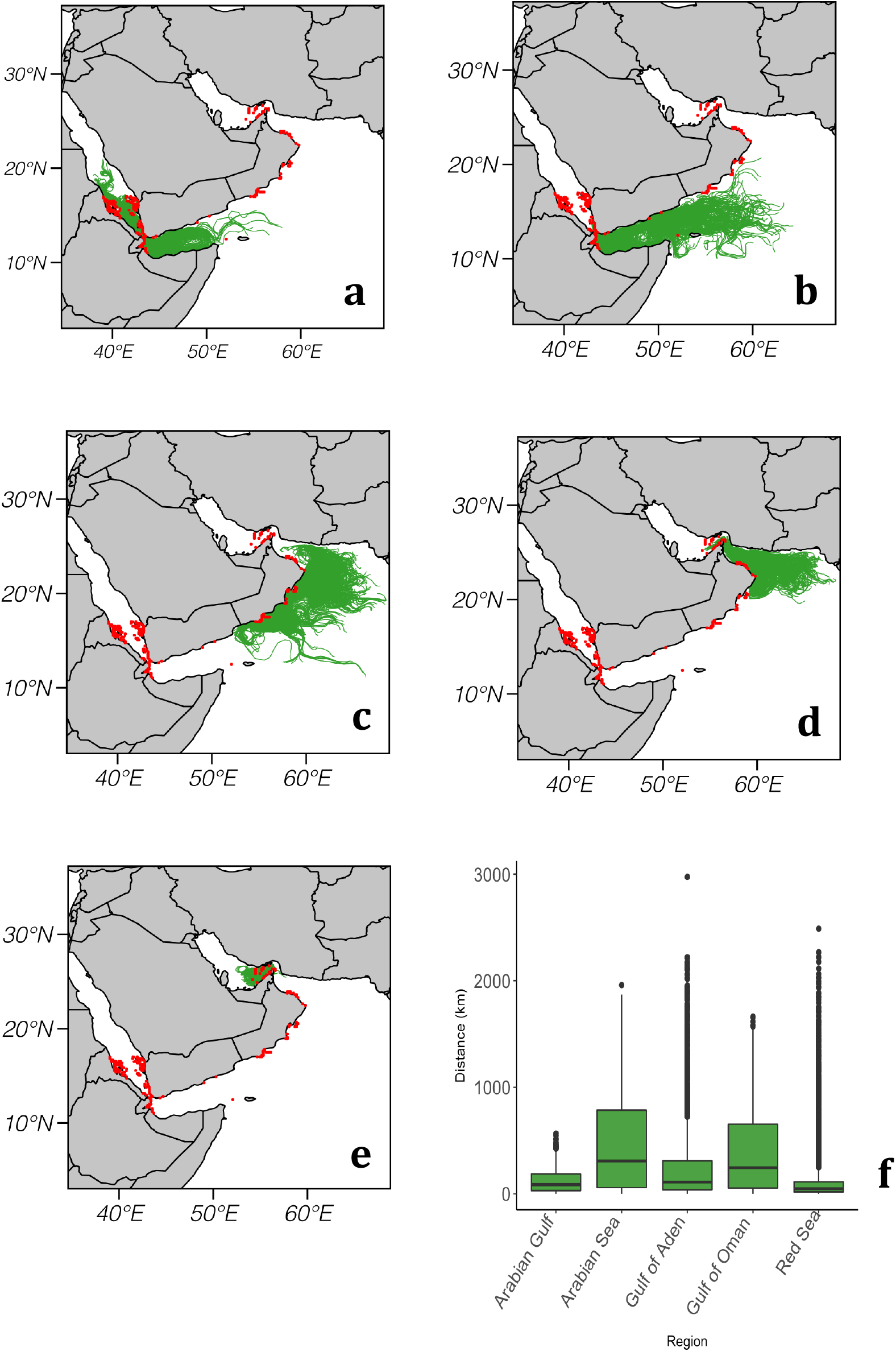
Modeled dispersal paths of epipelagic virtual larvae with larval duration of 40 days released in the summertime from (a) the Red Sea, (b) Gulf of Aden, (c) Arabian Sea, (d) Sea of Oman and (e) Arabian Gulf. For sake of visualization one ne larva was released from each habitat cell (total of 181) per day in 2015. The distances travelled by the larvae are represented in (f).

In the Arabian Sea, surface circulation exhibited a strong seasonal cycle. During the summer, larvae released along the Omani coast between 54.6° E and 56.1° E, either moved eastward alongshore toward the Sea of Oman or sharply turned toward the open ocean. Whereas, larvae released between 57.8° E and 59° E were exclusively transported alongshore toward the Sea of Oman (Figure S5). In the winter, even though the larvae along the entire Omani coast moved toward offshore, on average they travelled shorter distances compared to the summer (Figure 5f, 6f).

On the east side of the Arabian Peninsula, the temporal variability of both the direction and the distance travelled by the larvae was less evident. In the Sea of Oman, larvae released from the eastern Omani coast tended to move southward and few larvae reached the Gulf. However, those larvae originating on the top of the Sea of Oman were able to pass through the Strait of Hormuz, though they did not travel far away. In the Gulf the patterns of dispersal observed in both seasons were quite similar, larvae tended not to move far away and hence most of them remained in the Gulf.

## DISCUSSION

Measuring the strength of barriers has been a great challenge to understand phylo- and bio-geographic patterns (Treml et al., 2015). In our study, a series of individual-based simulations were performed to assess the permeability of larvae of 72 hypothetical taxa through three putative barriers. This ‘multitaxa’ comparison demonstrated how biological-physical interactions determine the success of propagules being transported through the Bab el-Mandeb Strait and the Strait of Hormuz, as well as of those being self-replenished off Oman. Here, we hypothesized that coupling hydrodynamic, seascape features and larval traits, potentially contribute to the distribution of genetic lineages since the connectivity rate, as shown in this study, represents a proxy of gene flow. Therefore, we assumed that the absence, strong reduction or asymmetry of larval exchange through the putative barriers affects the phylogeographic pattern around the Arabian Peninsula.

Given the aim of our study, we took advantages of a Lagrangian three-dimensional approach to assess the effects of the hydrographic variability conditions (e.g., extreme events, perturbation, instability) on larval dispersal. Langragian models, although more realistic, have computational requirements that make impossible to release a real number of larvae per species. Alternatively, Eulerian advection-diffusion methods, though are not spatially realistic, have a large impact on dispersal kernel and hence are suitable in quantifying evolutionarily significant tails in evolutionary connectivity studies (Treml et al., 2012).

Our results showed that the hardest putative barrier was that positioned off Oman, with scenarios exhibiting weak or none connectivity between the Arabian Sea and its adjacent areas. This result corroborates with previous population genetic studies. For example, phylogeographic investigations on fish species such as *Scomberomorus commerson* (van Herwerden et al., 2006); *Cephalopholis hemistiktos* (Priest et al., 2016) and *Pomacanthus maculosus* (Torquato et al., 2019), used different genetic markers and found a significant discontinuity positioned in southern Oman. Moreover, in our models the strength of this barrier increased when the taxa are spawning during the summer monsoon. And this prediction is in accordance with the gonado-somatic index (GSI) values for the three species mentioned above, i.e., *S. commerson* (Kaymaram et al., 2010); *C. hemistiktos* (Priest et al., 2016) and *P. maculosus* (Grandcourt et al., 2010), which indicate that they spawn mainly in this season. By contrast, among six other exploited finfish species inhabiting the Omani coast, only one (*Epinephelus diacanthus*) also revealed higher GSI values during the onset of the summer monsoon period (McIlwain et al., 2006).

The main hypotheses to explain faunal and genetic differences between both sides of the peninsula usually involve seascape features and/or ocean circulation off Oman (Torquato et al. 2019). The seascape is characterized by to coral colonies in southern Omani coast being weakly developed and represented by a reduced number of species (Burt et al., 2016; Sheppard & Salm, 1988), such that the lack of coral-habitat creates an unbridgeable gap for coral-dependent species (Priest et al., 2016). In turn, the ocean circulation explanation relies on studies involving small pelagic fishes, which suggest that upwelling systems hindrance connectivity by displacing larvae offshore due to Ekman transport (Parrish et al., 1981; but see Morgan et al., 2012). The Benguela upwelling system, for example, acts as a barrier to some species of phytoplankton, copepods and pelagic fishes between North and South (Lett et al., 2007).

Regarding the straits, our biophysical models showed an asymmetric movement thought both the Bab el-Mandeb Strait and the Strait of Hormuz. The strength of the first is in accordance with the seasonal water exchange pattern between the Red Sea and Gulf of Oman. The water flow through the Bab el-Mandeb Strait changes from a two-layer surface flow in the winter to a three-layer flow in the summer (Smeed, 2004). Thus, vertical migration combined with seasonal spawning play a critical role in the connectivity (Paris & Cowen, 2004), by positing the larvae in one or another prevailing current. For example, although larvae released from the Gulf of Aden always reached the Red Sea, larvae released from the second and that did not exhibit vertical migration, increased their retention significantly within this sea during the winter. Although ichthyoplankton surveys in the Red Sea indicated that the vast majority of fish taxa inhabiting the Red Sea spawn mainly during spring and summer (e.g., *Amphiprion bicinctus*) a few species actually spawn in the winter (El-Regal, 2013) and hence their larvae are subject to being retained regardless their PLD.

Investigations performed hitherto have not showed genetic discontinuities of species through the Bab el-Mandeb Strait, including fishes *A. bicinctus* (Saenz-Agudelo et al., 2015); *C. hemistiktos* (Priest et al. 2016); *P. maculosus*(Torquato et al., 201); *Chaetodon* spp. (DiBattista et al., 2020) and sea anemones (Emms et al., 2019). In fact, the strength of the barrier has been mainly debated based on the species distribution studies. Kemp (1998) suggested that although the Bab el-Mandeb Strait was a site of a significant Pleistocene vicariance event, it does not act as a present-day barrier. According to the author, the paucity of information about the reef fish assemblage inhabiting the southern Red Sea and the adjacent Gulf of Aden is the main reason for the hypothesis that the strait acts as a present-day barrier.

In the Strait of Hormuz, in turn, larvae exhibiting short PLD, experiencing high mortality, and especially those performing vertical migration, displayed great chance of not crossing the strait when released from the Sea of Oman. This combination of biological traits is likely observed in the study area, since the extreme thermal condition in the region may shorten PLDs, or even induce larvae to experience high mortality rates (Munday et al., 2009). Therefore, the consequent reduction in spatial scale of connectivity due to physical-biological interactions may be underling genetic differentiation between the adjacent populations, by reducing gene flow to levels that are unable to overcome local differentiations.

There are few evidences of the Strait of Hormuz acting as a barrier for coral-associated fauna. Weak genetic discontinuity between the Gulf and the adjacent Sea of Oman has been shown for a coral species *Platygyra daedalea* (Howells et al., 2016), which spawn mainly from February to May (Bauman et al., 2011), and a sea urchin *Echinometra* sp. (Ketchum et al., 2020), whose PLD within the genus vary from 18 to 30 days (McClanahan & Muthiga, 2007). On the other hand, population genetics studies showed that for reef fish species, such as *C. hemistiktos* (Priest et al., 2016) and *P. maculosus* (Torquato et al., 2019), the Gulf and the Sea of Oman represent a single phylogeographic province.

## Conclusion

Our biophysical models complement the existing research on comparative fish population genetics by providing a snapshot of the present-day seascape permeability around the Arabian Peninsula. The comparative and cross-taxon models could identify key biological traits and biophysical interactions that limit the transport of ichthyoplankton in our study area. The predictions presented here serve as testable hypotheses for future studies on population genetics, especially demographic models focusing on symmetry of gene flow, and fish larval distributions around the Arabian Peninsula.

## Acknowledgements

The author F.T. is supported by a CNPq/Brazil fellowship through the programme Science Without Borders (Proc. 232875/2014-6). We also thank Dr Robert Cowen (Oregon University) and Dr Luiz Rocha (California Academy of Sciences) for their helpful comments.

## BIOSKETCH

**Felipe Torquato** is interested in the origin and distribution of biodiversity at all ecological levels. This work represents a component of his PhD work at the Natural History Museum of Denmark under the supervision of Peter R. Møller.

## Author contributions

FT and PRM conceived the idea; FT designed and ran the biophysical models, and led the writing with assistance from PRM.

## References

Ayre, D. J, Minchinton, T. E., & Perrin, C. (2009). Does life history predict past and current connectivity for rocky intertidal invertebrates across a marine biogeographic barrier? Molecular Ecology, 18, 1887–1903.

Bauman, A. G., Baird, A. H., & Cavalcante, G. H. (2011). Coral reproduction in the world’s warmest reefs: southern Persian Gulf (Dubai, United Arab Emirates). Coral Reefs, 30, 405–413.

Baums, L. B., Paris, C. B., & Chérubin, L. M. (2006). A bio-oceanographic filter to larval dispersal in a reef-building coral. Limnology and Oceanography, 51, 1969–1981.

Berumen, M. L., DiBattista, J. D., & Rocha, L. A. (2017). Introduction to virtual issue on Red Sea and Western Indian Ocean biogeography. Journal of Biogeography, 44, 1923–1926.

Bowen, B. W., Gaither, M. R., DiBattista, J. D., Iaccheia, M., Andrews, K. R., Grant, W. S., Toonen, R. J., & Briggs, J. C. (2016). Comparative phylogeography of the ocean planet. Proceedings of the National Academy of Sciences, 113, 7962–7969.

Burt, J. A., Feary, D. A., Bauman, A. G., Usseglio, P., Cavalcante, G. H., Sale, & P. F. (2011). Biogeographic patterns of reef fish community structure in the northeastern Arabian Peninsula. ICES Journal of Marine Science, 68, 1875–1883.

Burt, J. A., Coles, S., van Lavieren, H., Taylor, O., Looker, E., & Samimi-Namin, K. (2016). Oman’s coral reefs: A unique ecosystem challenged by natural and man- related stresses and in need of conservation. Marine Pollution Bulletin, 105, 498–506.

Cowen, R. K., Lwiza, K. M. M., Sponaugle, S., Paris, C. B., & Olson, D. B. (2000). Connectivity of Marine Populations: Open or Closed? Science, 287, 857–859.

Cowen, R. K. (2002). Larval dispersal and retention and consequences for population connectivity. In Sale, P. F. (Ed.), Coral reef fishes: dynamics and diversity in a complex ecosystem (pp. 149–170). Academic Press.

Cowen, R. K., & Sponaugle, S. (2009). Larval Dispersal and Marine Population Connectivity. Annual Review of Marine Science,1, 443–466.

Cribari-Neto, F., & Zeileis, A. (2010). Beta Regression in R. Journal of Statistical Software, 34, 1–24.

Cutler, A. N., & Swallow, J. C. (1984). Surface currents of the Indian Ocean (to 25S, 100E): compiled from historical data archived by the Meteorological Office, Institute of Oceanographic Sciences.

DiBattista, J. D., Berumen, M. L., Gaither, M. R., Rocha, L. A., Eble, J. A., Choat, J. H., … Bowen, B. W. (2013). After continents divide: comparative phylogeography of reef fishes from the Red Sea and Indian Ocean. Journal of Biogeography, 40, 1170–1181.

DiBattista, J. D., Waldrop, E., Rocha, L. A., Craig, M. T., Berumen, M. L., & Bowen, B. W. (2015). Blinded by the bright: A lack of congruence between colour morphs, phylogeography and taxonomy for a cosmopolitan Indo‐Pacific butterflyfish, Chaetodon auriga. Journal of Biogeography, 42, 1919–1929.

DiBattista, J. D., Roberts, M. B., Bouwmeester, J., Bowen, B. W., Coker, D. J., Lozano-Cortes, D. F., … Berumen, M. L. (2016). A review of contemporary patterns of endemism for shallow water reef fauna in the Red Sea. Journal of Biogeography, 43, 423–439.

DiBattista, J. D., Choat, J. H., Gaither, M. R., Hobbs, J. A., Lozano-Cortes, D. F., Myers, R. F., … Berumen, M. L. (2016). On the origin of endemic species in the Red Sea. Journal of Biogeography, 43, 13–30.

DiBattista, J. D., Gaither, M. R., Hobbs, J. A., Saenz-Agudelo, P., Piatek, M. J., Bowen, B. W., … Berumen, M. L. (2017). Comparative phylogeography of reef fishes from the Gulf of Aden to the Arabian Sea reveals two cryptic lineages. Coral Reefs, 36, 625–638.

DiBattista, J. D., Saenz-Agudelo, P., Piatek, M. J., Cagua, E. F., Bowen, B. W., Choat, J. H., … McIlwain, J. H. (2020). Population genomic response to geographic gradients by widespread and endemic fishes of the Arabian Peninsula. Ecology and Evolution, 10, 4314–4330.

El-Regal, M. A. (2013). Spawning seasons, spawning grounds and nursery grounds of some Red Sea fishes. The Global Journal of Fisheries and Aqua, 6, 105–125.

Elliott, A. J., & Savidge, G. (1990). Some features of the upwelling off Oman. Journal of Marine Research, 48, 319–333.

Ferrari, S., & Cribari-Neto, F. (2004). Beta Regression for Modelling Rates and Proportions. Journal of Applied Statistics, 31, 799–815.

Galarza, J. A., Carreras-Carbonell, J., Macpherson, E., Pascual, M., Roques, S., Turner, G. F., & Rico, C. (2009). The influence of oceanographic fronts and early-life-history traits on connectivity among littoral fish species. Proceedings of the National Academy of Sciences, 106, 1473–1478.

Grandcourt, E., Al Abdessalaam, T. Z., Francis, F., & Al Shamsi, A. (2010). Age‐based life history parameters and status assessments of by‐catch species (Lethrinus borbonicus, Lethrinus microdon, Pomacanthus maculosus and Scolopsis taeniatus) in the southern Arabian Gulf. Journal of Applied Ichthyology, 26, 381–389.

Holstein, D. M., Paris, C. B., & Mumby, P. J. (2014). Consistency and inconsistency in multispecies population network dynamics of coral reef ecosystems. Marine Ecology Progress Series, 499, 1–18.

Howells, E. J., Abrego D., Meyer E., Kirk N. L. & Burt J. A. (2016). Host adaptation and unexpected symbiont partners enable reef-building corals to tolerate extreme temperatures. Global Change Biology, 22, 2702–2714.

Irisson, J. O., Paris, C. B., Guigand, C., & Planes, S. (2010). Vertical distribution and ontogenetic ‘‘migration’’ in coral reef fish larvae. Limnology and Oceanography, 55, 909–919.

Kaymaram, F., Hossainy, S. A., Darvishi, M., Talebzadeh, S. A., & Sadeghi, M. S. (2010). Reproduction and spawning patterns of the Scomberomorus commerson in the Iranian coastal waters of the Persian Gulf & Oman Sea. Iranian Journal of Fisheries Sciences, 9, 233–244.

Kemp, J. M. (1998). Zoogeography of the coral reef fishes of the Socotra Archipelago. Journal of Biogeography, 25, 919–933.

Ketchum, R. N., Smith, E. G., DeBiasse, M. B., Vaughan, G. O., McParland, D., Leach, W. B., … Reitzel, A. M. (2020). Population genomic analyses of the sea urchin Echinometra sp. EZ across an extreme environmental gradient. Genome biology and evolution.

Lett, C., Veitch, J., van der Lingen, C. D., & Hutchings, L. (2007). Assessment of an environmental barrier to transport of ichthyoplankton from the southern to the northern Benguela ecosystems. Marine Ecology Progress Series, 347, 247–259.

Lindeman, K. C., Richards, W. J., Lyczkowski-Shultz, J., Drass, D. M., Paris, C. B., Leis, J. M., Lara, M., & Comyns, B. H. (2005). Lutjanidae: snappers. In Richards, W. J. (Ed.), Early stages of Atlantic fishes (pp. 1549–1586). CRC Press.

McClanahan TR, & Muthiga N. A. (2007). Chapter 15 Ecology of Echinometra. In: Lawrence JM, editor. Developments in Aquaculture and Fisheries Science: Elsevier. p. 297–317.

McIlwain, J., Hermosa, G. V., Claereboudt, M., Al-Oufi, H. S., & Al-Awi, M. (2006). Spawning and reproductive patterns of six exploited finfish species from the Arabian Sea, Sultanate of Oman. Journal of Applied Ichthyology, 22, 167–176.

Morgan, S. G., Fisher, J. L., McAfee, S. T., Largier, J. L., & Halle, C. M. (2012). Limited recruitment during relaxation events: larval advection and behavior in an upwelling system. Limnology and Oceanography, 57, 457–470.

Munday, P. L., Leis, J. M., Lough, J. M., Paris, C. B., Kingsford, M. J., Berumen, M. L., & Lambrechts, J. (2009). Climate change and coral reef connectivity. Coral reefs, 28, 379–395.

Nanninga, G. B., Saenz-Agudelo, P., Manica, A., & Berumen, M. L. (2014). Environmental gradients predict the genetic population structure of a coral reef fish in the Red Sea. Molecular Ecology, 23, 591–602.

Paris, C. B., & Cowen, R. K. (2004). Direct evidence of a biophysical retention mechanism for coral reef fish larvae. Limnology and Oceanography, 49, 1964–1979.

Paris, C. B., Helgers, J., Sebille, E., & Srinivasan, A. (2013). Connectivity Modeling System: A probabilistic modeling tool for the multi-scale tracking of biotic and abiotic variability in the ocean. Environmental Modelling & Software, 42, 47–54.

Parrish, H. R., Nelson, C. S., & Bakun, A. (1981). Transport mechanisms and reproductive success of fishes in the California current. Biological Oceanography, 1, 175–203.

Priest, M. A., DiBattista, J. D., McIlwain, J. L., Taylor, B. M., Hussey, N. E., & Berumen, M. L. (2016). A bridge too far: dispersal barriers and cryptic speciation in an Arabian Peninsula grouper (Cephalopholis hemistiktos). Journal of Biogeography, 43, 820–832.

Rocha, L. A. (2003). Patterns of distribution and processes of speciation in Brazilian reef fishes. Journal of Biogeography, 30, 1161–1171.

Saenz-Agudelo, P., DiBattista, J. D., Piatek, M. J., Gaither, M. R., Harrison, H. B., Nanninga, G. B., & Berumen, M. L. (2015). Seascape genetics along environmental gradients in the Arabian Peninsula: insights from ddRAD sequencing of anemonefishes. Molecular Ecology, 24, 6241–6255.

Sheppard, C. R. C., & Salm, R. V. (1988). Reef and coral communities of Oman, with a description of a new coral species (Order Scleractinia, genus Acanthastrea). Journal of Natural History, 22, 263–279.

Shetye, S. R., Gouveia, A. D., & Shenoi, S. C. (1994). Circulation and water masses of the Arabian Sea. Proceedings of the Indian Academy of Sciences, 103, 107–123.

Smeed, D. A. (2004). Exchange through the Bab el Mandab. Deep-Sea Research II, 51, 455–474.

Thresher, R. E., & Brothers, E. B. (1985). Reproductive ecology and biogeography of Indo-West Pacific angelfishes. Evolution. 39: 878–887.

Torquato, F. O., Range, P., Ben-Hamadou, R., Sigsgaard, E. E., Thomsen, P. F., Riera, R., … Møller, P. R. (2019). Consequences of marine barriers on genetic diversity of the coral-specialist yellowbar angelfish from the western Indian Ocean. Ecology and Evolution, 9, 11215–11226.

Treml, E. A., Roberts, J. J., Chao, Y., Halpin, P. N., Possingham, H. P. & Riginos, C. (2012). Reproductive Output and Duration of the Pelagic Larval Stage Determine Seascape-Wide Connectivity of Marine Populations. Integrative and Comparative Biology, 52, 525–537.

Treml, E. A., Roberts, J., Halpin, P. N., Possingham, H. P., & Riginos, C. (2015). The emergent geography of biophysical dispersal barriers across the Indo-West Pacific. Diversity and Distributions, 21, 465–476.

UNEP-WCMC, WorldFish Centre, WRI, TNC (2010). Global distribution of warm-water coral reefs, compiled from multiple sources including the Millennium Coral Reef Mapping Project. Version 3.0. Includes contributions from IMaRS-USF and IRD (2005), IMaRS-USF (2005) and Spalding et al. (2001). UN Environment World Conservation Monitoring Centre, Cambridge/UK. URL: http://data.unep-wcmc.org/datasets/1.

van Herwerden, L., McIlwain, J., Al-Oufi, H., Al-Amry, W., & Reyes, A. (2006). Development and application of microsatellite markers for Scomberomorus commerson (Perciformes; Teleostei) to a population genetic study of Arabian Peninsula stocks. Fisheries Research, 79, 258–266.

Zuur, A. F., Ieno, E. N., & Elphick, C. S. (2010). A protocol for data exploration to avoid common statistical problems. Methods in Ecology and Evolution, 1, 3–14.

