## Supplemental Information for "Physical-biological interactions underlying the phylogeographic patterns of coral-dependent fishes around the Arabian Peninsula"

### Appendix S1.

##### Assessing model saturation:

##### Methods

Sensitivity analyses of some parameters can potentially improve the accuracy of the model (North et al., 2009). Here, the releasing frequency was computed to saturate all possibilities of dispersal based on the connectivity matrix. To determine the release frequency, we conducted a numerical experiment over five months in 2015 (January, March, June, September and December). In this experiment particles were release from the 181 polygons, and the number of released particles remained constant (one hundred) while the releasing frequency varied from 1.5 hours (baseline) to 2 days (1.5 h, 3 h, 6 h, 12 h, 24 h, 48 h).

All values were extracted from the average matrix, vectored and used to calculate the fraction of unexplained variance (FUV) (Simons et al., 2013). To do so, the simulation that released 100 particles/1.5h over the five months was used as the baseline and compared to their peers in a linear model. The coefficient of determination ($r^2$) was then calculated between each baseline/increment pairing. A 0.05 threshold FUV variance was used to define the point where variance in FUV was minimal (Ross et al., 2016; Simons et al., 2013).

The same experiment was conducted to determine the value of horizontal diffusion by comparing three values, namely: 5 $m.s^{-1}$, 50 $m.s^{-1}$ and 100 $m.s^{-1}$.

**Results**

Release frequency showed no significant differences whether 100 particles were released every 1.5 or 24 hours. The second experiment, in turn showed that 50 m.s$^{-1}$ was significantly different compared to 5 m.s$^{-1}$, but not different compared to 100 m.s$^{-1}$.

#### Release Frequency

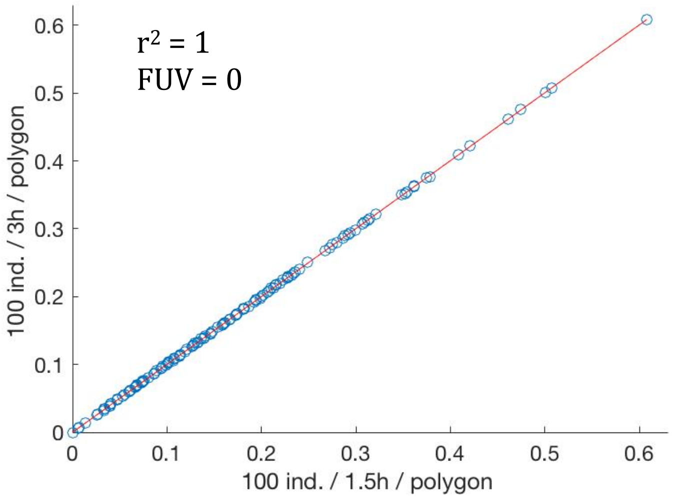


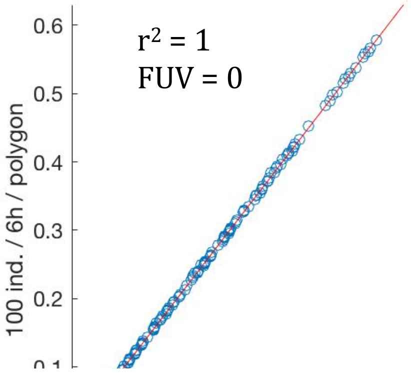


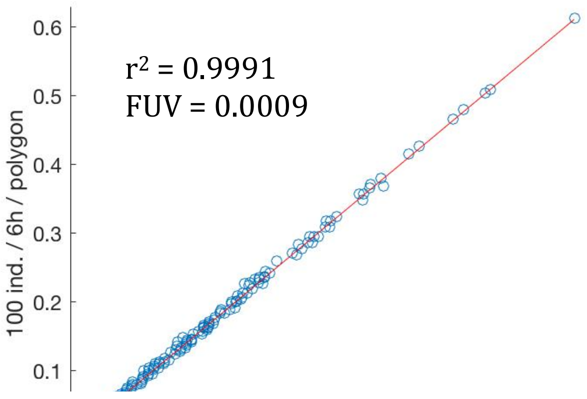

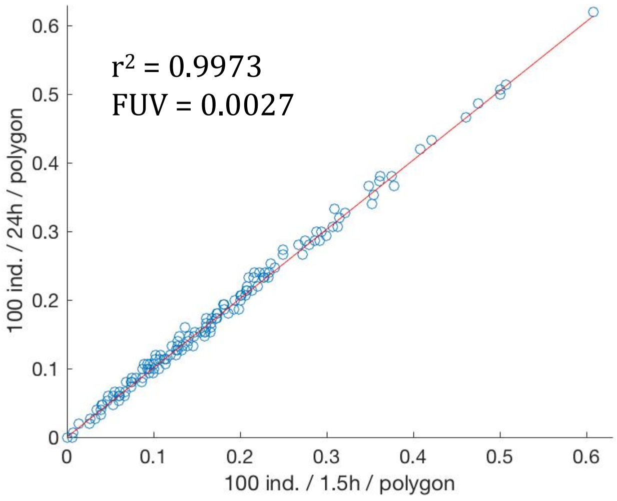


**Horizontal Diffusion**

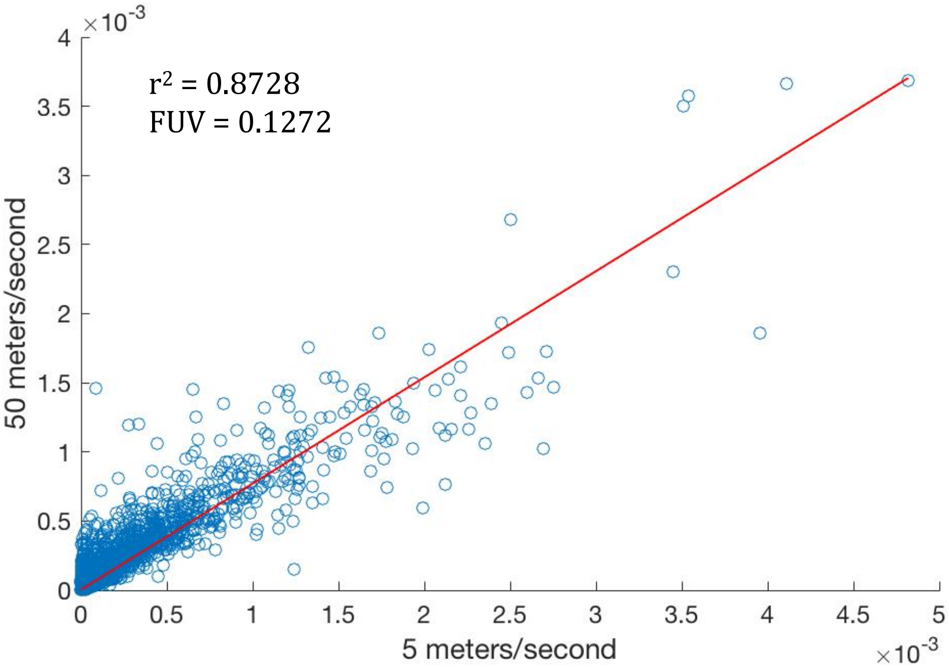


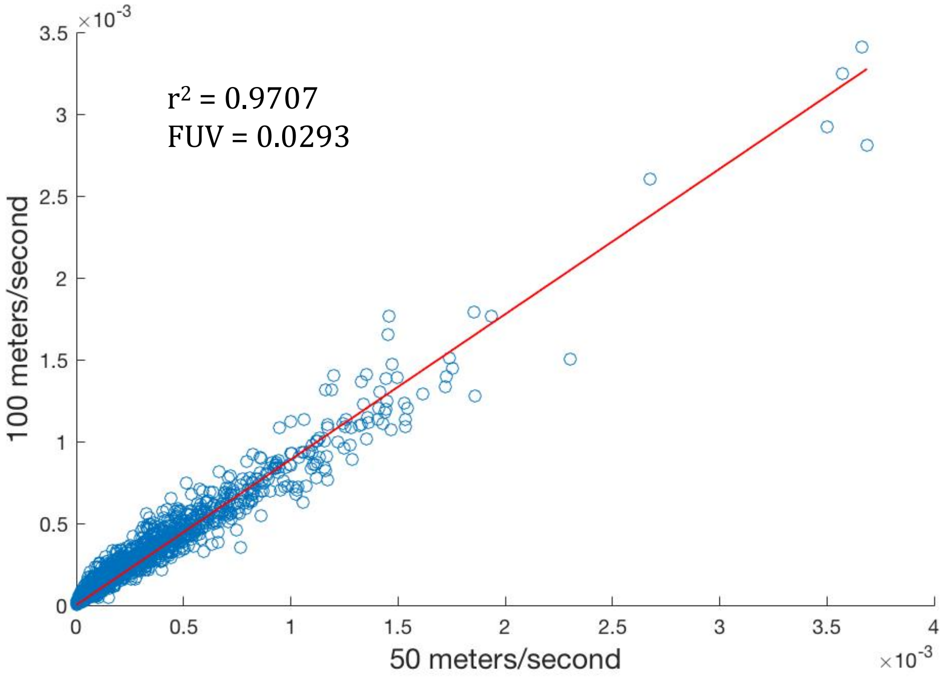


### TABLES

**Table S1.** Summary statistics of the tested beta regression models, where (*) highlights significant explanatory variables at a level of 5% of significance.

| Bab-el-Mandeb Strait (Red Sea : Gulf of Aden) | | | | | |
|---|---|---|---|---|---|
| | **Estimate** | **Std. Error** | **z-value** | **p-value** | **R-squared** |
| Intercept | -3.96972 | 0.07015 | -56.591 | < 2e-16 *** | 0.8189 |
| PLD: 30 | 0.20271 | 0.05761 | 3.518 | 0.000434 *** | |
| PLD: 40 | 0.32290 | 0.05627 | 5.739 | 9.54e-09 *** | |
| Mortality: low | 0.54020 | 0.04663 | 11.585 | < 2e-16 *** | |
| Vertical migration | 0.38233 | 0.04593 | 8.324 | < 2e-16 *** | |
| Spawning: winter | -0.37049 | 0.06022 | -6.152 | 7.66e-10 *** | |
| Spawning: summer | 0.23742 | 0.05226 | 4.543 | 5.55e-06 *** | |
| Reproductivity: high | 0.01098 | 0.04525 | 0.243 | 0.808243 | |
| **Bab-el-Mandeb Strait (Gulf of Aden : Red Sea)** | | | | | |
| Intercept | -1.983452 | 0.070345 | -28.196 | < 2e-16 *** | 0.8605 |
| PLD: 30 | 0.280960 | 0.061274 | 4.585 | 4.53e-06 *** | |
| PLD: 40 | 0.408091 | 0.060225 | 6.776 | 1.23e-11 *** | |
| Mortality: low | 0.628795 | 0.049457 | 12.714 | < 2e-16 *** | |
| Vertical migration | -0.84981 | 0.050465 | -16.840 | < 2e-16 *** | |
| Spawning: winter | 0.217171 | 0.056316 | 3.856 | 0.000115 *** | |
| Spawning: summer | -0.416469 | 0.062810 | -6.631 | 3.34e-11 *** | |
| Reproductivity: high | 0.004908 | 0.048346 | 0.102 | 0.919138 | |

**Table S1.** Cont.

| Strait of Hormuz (Arabian Gulf : Gulf of Oman) | | | | | |
|---|---|---|---|---|---|
| | **Estimate** | **Std. Error** | **z-value** | **p-value** | **R-squared** |

| | | | | | |
|---|---|---|---|---|---|
| Intercept | -2.4377122 | 0.0742578 | -32.828 | < 2e-16 *** | 0.7908 |
| PLD: 30 | 0.2801397 | 0.0612423 | 4.574 | 4.78e-06 *** | |
| PLD: 40 | 0.3947315 | 0.0603014 | 6.546 | 5.91e-11 *** | |
| Mortality: low | 0.6300321 | 0.0494966 | 12.729 | < 2e-16 *** | |
| Vertical migration | 0.4250502 | 0.0488923 | 8.694 | < 2e-16 *** | |
| Spawning: winter | 0.0308873 | 0.0584299 | 0.529 | 0.5971 | |
| Spawning: summer | -0.1266759 | 0.0599479 | -2.113 | 0.0346 * | |
| Reproductivity: high | -0.0005956 | 0.0484149 | -0.012 | 0.9902 | |
| **Strait of Hormuz (Gulf of Oman : Arabian Gulf)** | | | | | |
| Intercept | -2.85255 | 0.02645 | -107.856 | <2e-16 *** | 0.9856 |
| PLD: 30 | 0.36306 | 0.02315 | 15.680 | <2e-16 *** | |
| PLD: 40 | 0.52537 | 0.02256 | 23.292 | <2e-16 *** | |
| Mortality: low | 0.58916 | 0.01833 | 32.142 | <2e-16 *** | |
| Vertical migration | -1.52844 | 0.02194 | -69.666 | <2e-16 *** | |
| Spawning: winter | 0.22018 | 0.02102 | 10.475 | <2e-16 *** | |
| Spawning: summer | -0.20977 | 0.02293 | -9.147 | <2e-16 *** | |
| Reproductivity: high | 0.02436 | 0.01780 | 1.369 | 0.171 | |

| Upwelling off Oman (Self-recruitment Arabian Sea) | | | | | |
|---|---|---|---|---|---|
| | **Estimate** | **Std. Error** | **z-value** | **p-value** | **R-squared** |
| Intercept | -1.0839437 | 0.0178398 | -60.760 | < 2e-16 *** | 0.9818 |
| PLD: 30 | -0.0811127 | 0.0151205 | -5.364 | 8.12e-08 *** | |
| PLD: 40 | -0.2013820 | 0.0152888 | -13.172 | < 2e-16 *** | |
| Mortality: low | 0.7439486 | 0.0125756 | 59.158 | < 2e-16 *** | |
| Vertical migration | 0.1018697 | 0.0124603 | 8.176 | 2.95e-16 *** | |
| Spawning: winter | 0.0007115 | 0.0151898 | 0.047 | 0.963 | |
| Spawning: summer | -0.0786474 | 0.0152944 | -5.142 | 2.71e-07 *** | |
| Reproducitivity: high | -0.0029511 | 0.0124581 | -0.237 | 0.813 | |

#### FIGURES

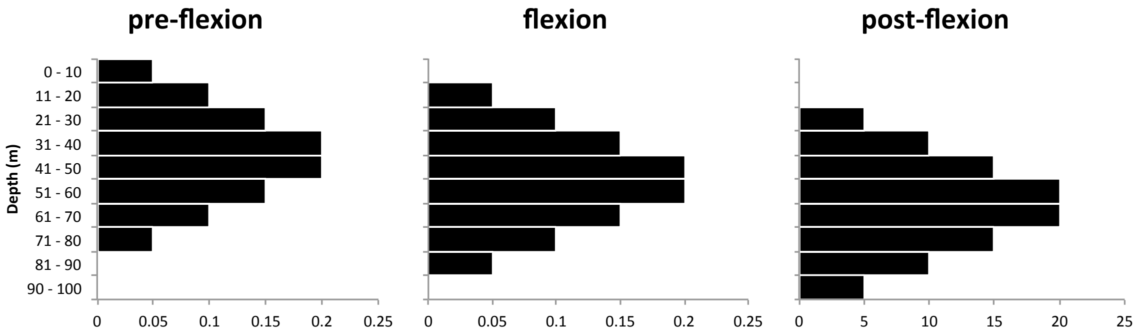


**Fig. S1.** Idealized vertical distribution of fish larvae at three ontogenetic stages.

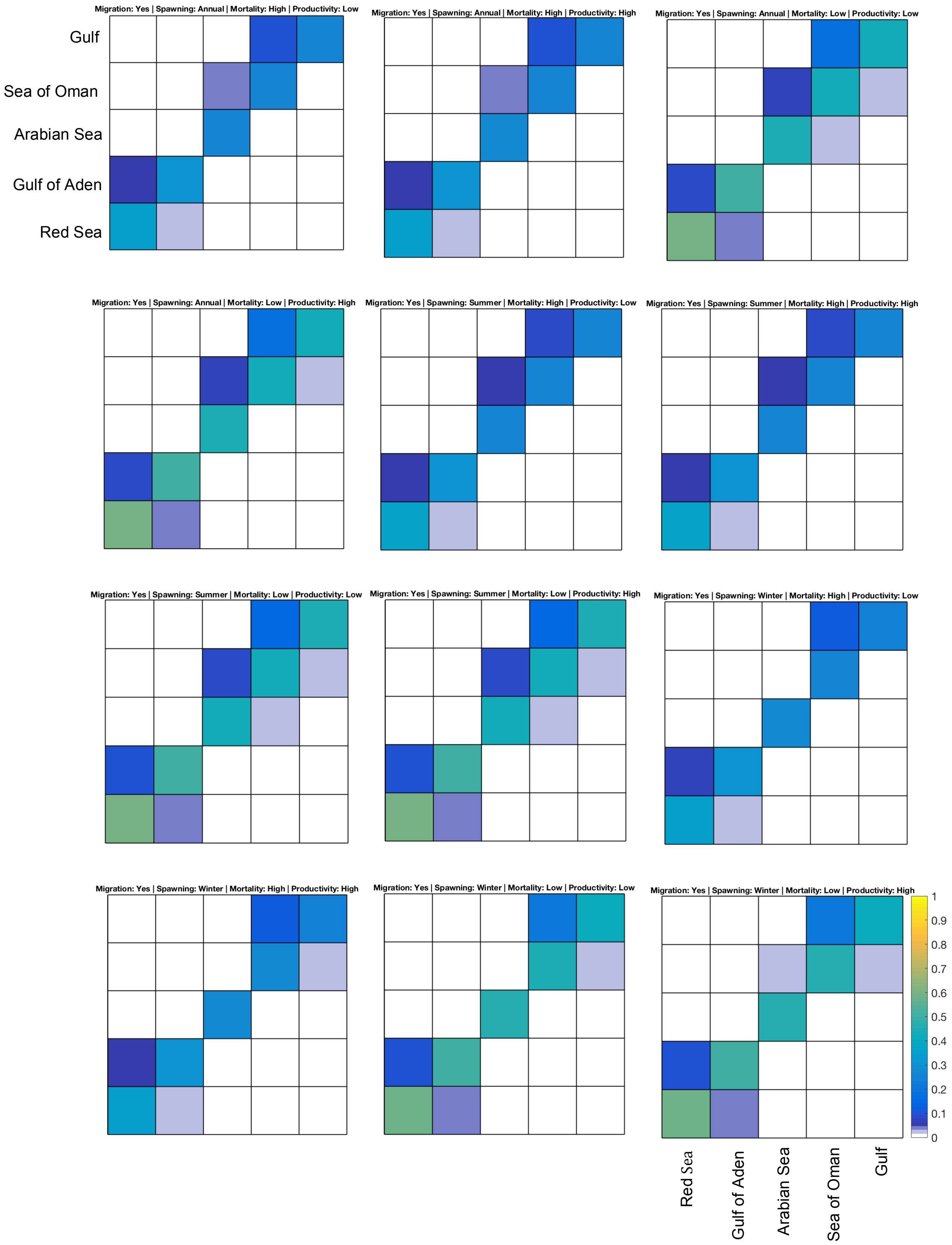


**Fig. S2.** Potential connectivity matrix with larval duration of 20 days. The colorbar shows the root square of the proportion of survived larvae exchanged between each release (y-axis) and settlement (x-axis) region.

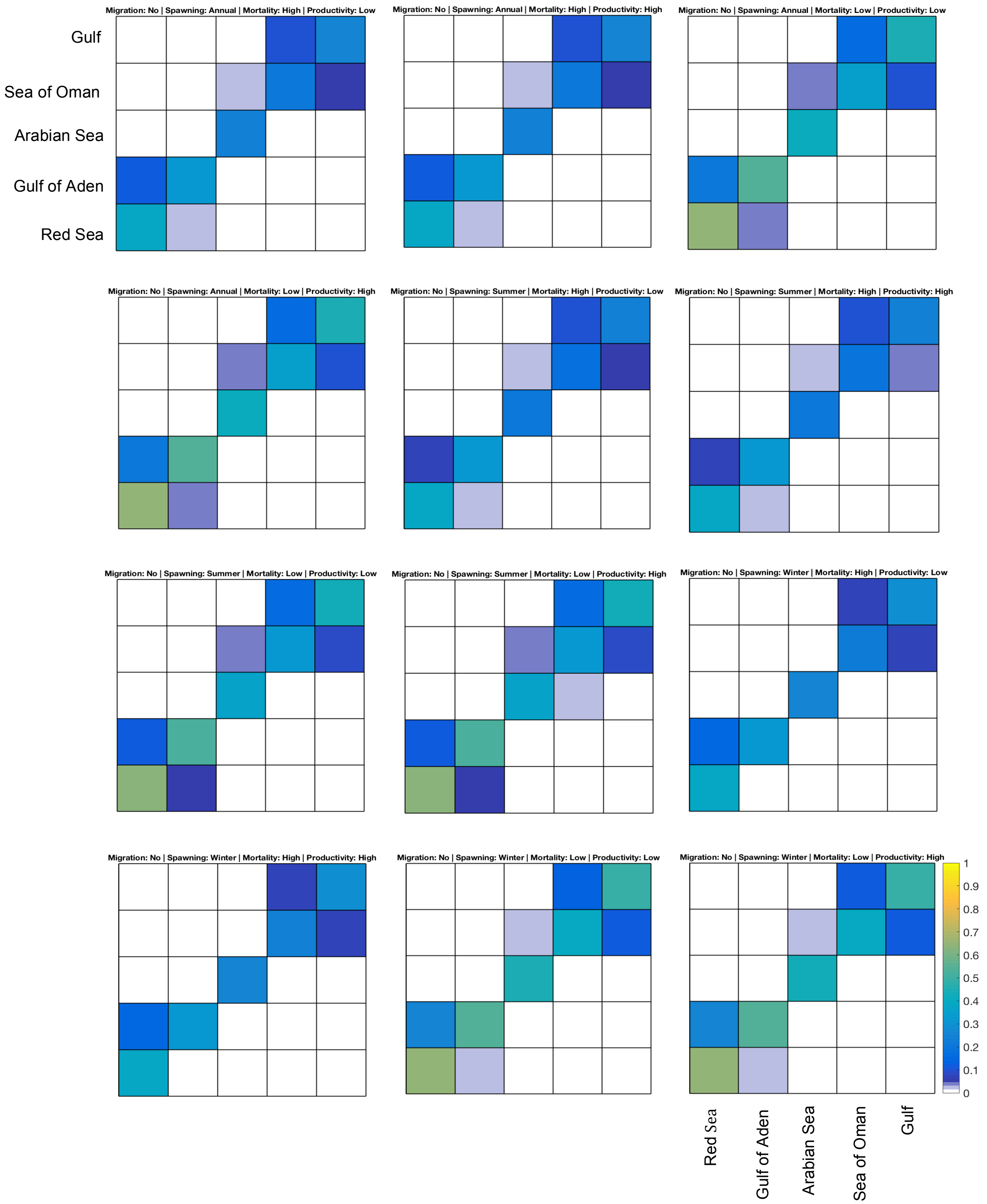


**Fig. S2 cont.** Potential connectivity matrix with larval duration of 20 days. The colorbar shows the root square of the proportion of survived larvae exchanged between each release (y-axis) and settlement (x-axis) region.

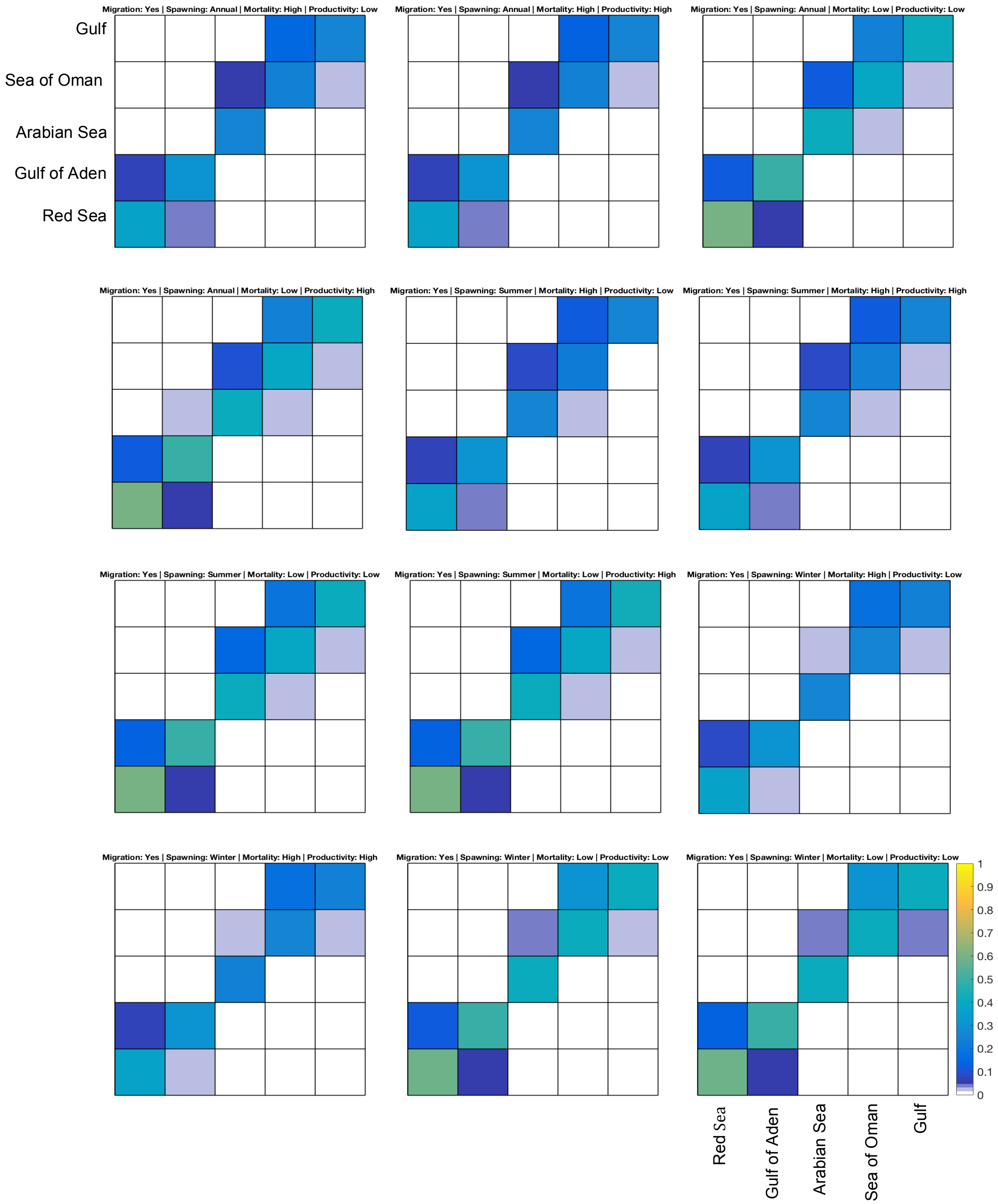


**Fig. S3.** Potential connectivity matrix with larval duration of 30 days. The colorbar shows the root square of the proportion of survived larvae exchanged between each release (y-axis) and settlement (x-axis) region.

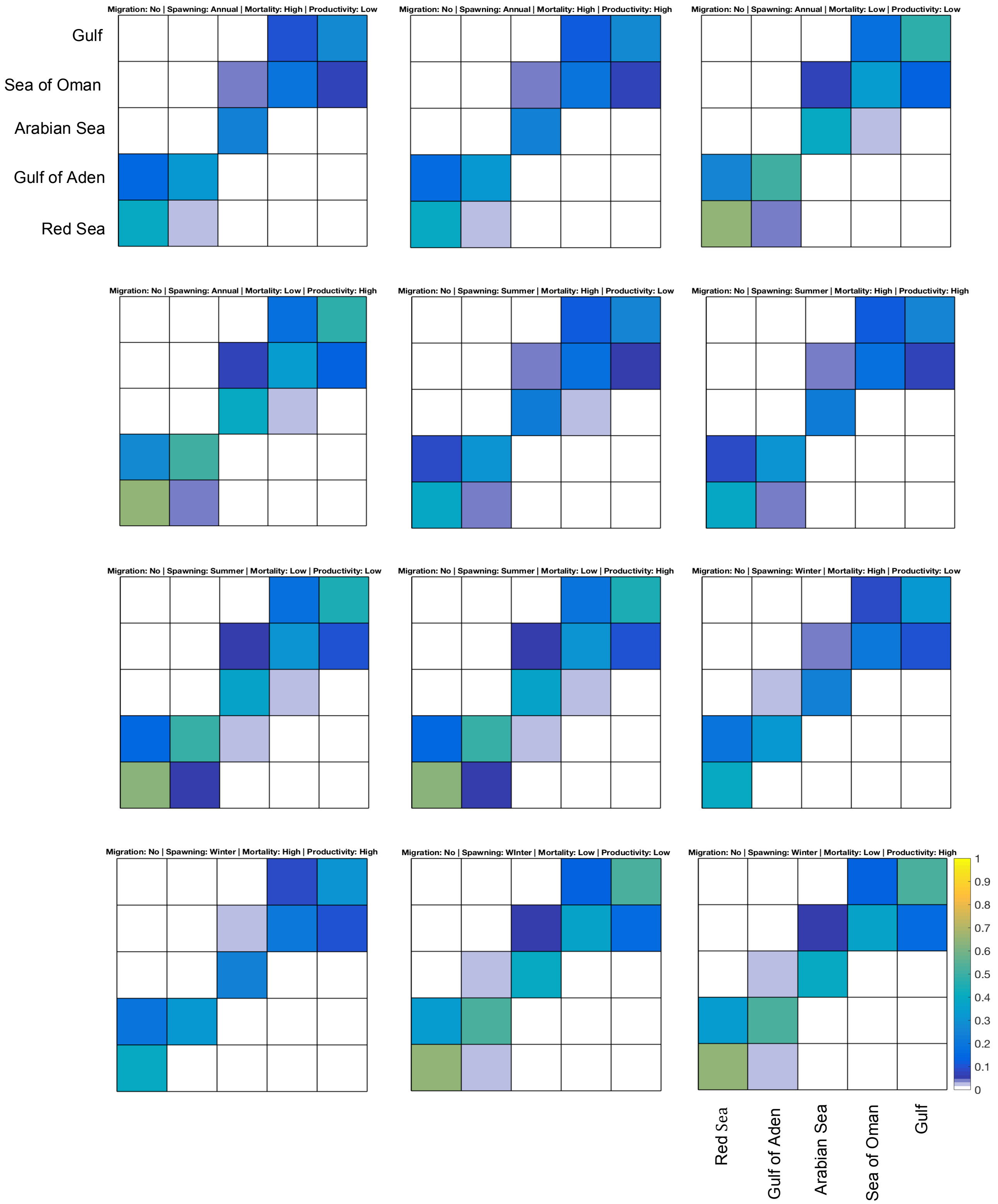


**Fig. S3 cont.** Potential connectivity matrix with larval duration of 30 days. The colorbar shows the root square of the proportion of survived larvae exchanged between each release (y-axis) and settlement (x-axis) region.

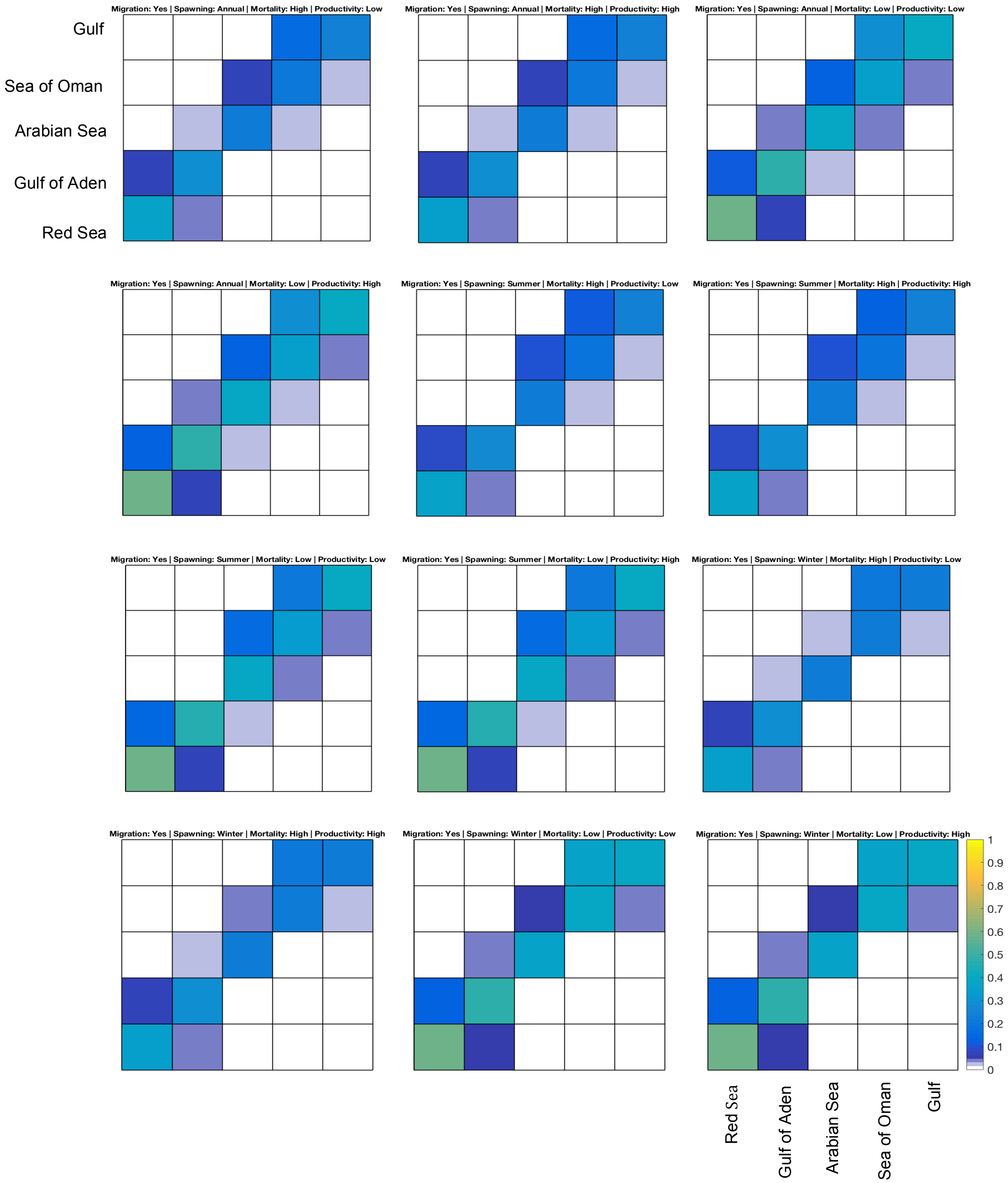


**Fig. S4.** Potential connectivity matrix with larval duration of 40 days. The colorbar shows the root square of the proportion of survived larvae exchanged between each release (y-axis) and settlement (x-axis) region.

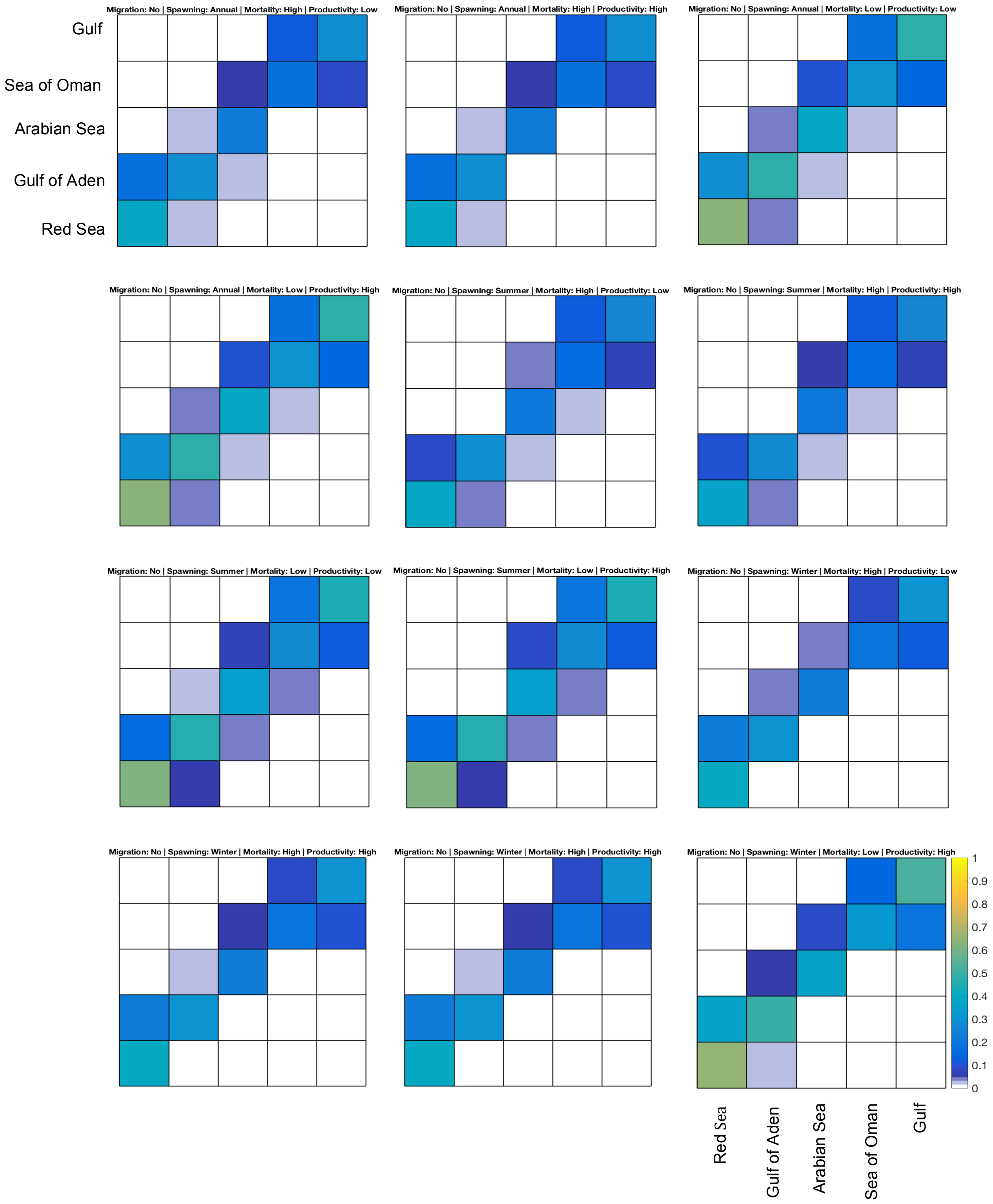


**Fig. S4 cont.** Potential connectivity matrix with larval duration of 40 days. The colorbar shows the root square of the proportion of survived larvae exchanged between each release (y-axis) and settlement (x-axis) region.

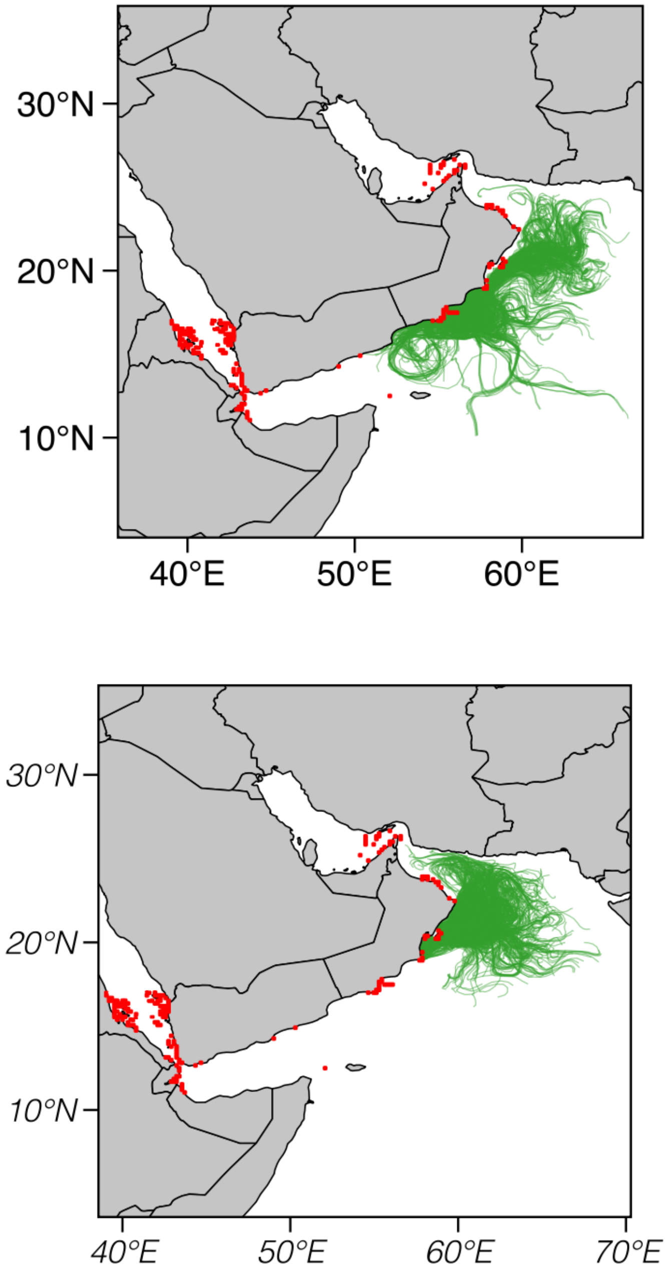


**Fig. S5.** Modeled dispersal paths of epipelagic virtual larvae with larval duration of 40 days released in the summertime from the Arabian Sea between (a) 54.6˚ E and 56.1˚ E , and (b) 57.8˚ E and 59˚ E.

**References**

North, E. W., Gallego, A., and Petitgas, P. 2009. Manual of recommended practices for modeling physical–biological interactions during fish early life, ICES Cooperative Research Report.

Ross, R. E., Nimmo-Smith, W. A. M., and Howell, K. L. 2016. Increasing the Depth of Current Understanding: Sensitivity Testing of Deep-Sea Larval Dispersal Models for Ecologists. PLoS ONE 11: e0161220.

Simons, R. D., Siegel, D. A., and Brown, K. S. 2013. Model sensitivity and robustness in the estimation of larval transport: A study of particle tracking parameters. J. Mar. Syst. 119–120: 19–29.
